# High-sensitivity detection of facial features on MRI brain scans with a convolutional network

**DOI:** 10.1101/2021.04.25.441373

**Authors:** Shashank Bansal, Avinash Kori, Wazeer Zulfikar, Joseph Wexler, Christopher J. Markiewicz, Franklin F. Feingold, Russell A. Poldrack, Oscar Esteban

## Abstract

Platforms and institutions that support MRI data sharing need to ensure that identifiable facial features are not present in shared images. Currently, this assessment requires manual effect as no auto-mated tools exist that can efficiently and accurately detect if an image has been “defaced”. The scarcity of publicly available data with preserved facial features, as well as the meager incentives to create such a cohort privately, have averted the development of face-detection models. Here, we introduce a framework to detect whether an input MRI brain scan has been defaced, with the ultimate goal of streamlining it within the submission protocols of MRI data archiving and sharing platforms. We present a binary (defaced/”nondefaced”) classifier based on a custom convolutional neural network architecture. We train the model on 980 de-faced MRI scans from 36 different studies that are publicly available at OpenNeuro.org. To overcome the unavailability of nondefaced examples, we augment the dataset by inpainting synthetic faces into each training image. We show the adequacy of such a data augmentation in a cross-validation evaluation. We demonstrate the performance estimated with cross-validation matches that of an evaluation on a held-out dataset (*N* =581) preserving real faces, and obtain accuracy/sensitivity/specificity scores of 0.978/0.983/0.972, respectively. Data augmentations are key to boosting the performance of models bounded by limited sample sizes and insufficient diversity. Our model contributes towards developing classifiers with *∼*100% sensitivity detecting faces, which is crucial to ensure that no identifiable data are inadvertently made public.

## 1 Introduction

Most existing and upcoming health-, and research-data privacy regulations [1, 2] are strengthening the protection of personally identifiable information (PII). In the case of neuroimaging, not only does such protection entail the removal of metadata that could directly and indirectly disclose PII (e.g., name, date of birth, date of acquisition, etc.), but also involves photographic PII that are present in some high-resolution, 3D magnetic resonance images (MRI) [2–4]. Therefore, before data can be transferred to uncertified systems for analysis and/or are made publicly available, they undergo a processing step typically referred to as “defacing” to remove data segments that would allow generating identifiable 3D models of a participant’s face [5]. Unsurprisingly, neuroimaging data management plans have globally adopted the anonymization of the data as the first, essential step of their workflow before data egresses the secure systems of the scanning facility. Moreover, data analysis infrastructure, data exchange services, and open-data sharing initiatives in the neuroimaging field must ensure that all their protocols and distribution do not have access to or unintendedly release PII.

Consequently, several automated tools such as `mri_deface` [5] and *PyDeface* [6] have been proposed and are widely adopted to identify and erase facial features from candidate MRI images (generally, high-resolution, anatomical MRI acquisitions). Typically, these methods align the participant’s image with a population average (also called “template”) where a mask enclosing all facial features to be removed has been manually defined. The estimated mapping between the template and participant’s image is then used to project the mask onto the target image, and all the voxels within the projected mask are zeroed [5]. Although these tools have demonstrated very high reliability, there are always underperforming instances, and facial features (nose, eyes, etc.) may partially or fully emerge unnoticed from the anonymization workflow.

Traditionally, before the processing and analysis, a visual quality control checkpoint is set up at the end of this data management workflow to capture these errors in defacing and ascertain other quality issues showcased by the data [7]. However, in the context of the data deluge that neuroimaging is witnessing, screening by one or more expert curators of every image belonging to sizeable datasets is impractical. To our knowledge, there are no tools that can automatically, and with very high sensitivity, determine that an MRI image has not been correctly defaced before publication, preventing the unintentional public disclosure of PII. Although the problem *a priori* seems an obvious task for learning a convolutional network, the very limited access (if any) to sufficiently large “nondefaced” datasets for training such models has steered the community’s effort in other directions. In addition to data unavailability, there is also the possibility that these models may internally learn and encode PII that can then be exploited to triangulate a participant’s identity. These limitations also pose stringent limits to the dataset’s diversity because composing an extensive and multi-site dataset of nondefaced data is just not feasible.

Here, we introduce an automated framework to detect nondefaced T1-weighted (T1w) images of the human brain. We leverage the OpenNeuro.org sharing initiative [8, RRID:SCR_005031] to compose a large, multi-site dataset of only defaced images. To overcome the unavailability of nondefaced examples and the risk of encoding PII within the model, we augment our dataset with synthetically generated data. To do so, we inpainted a realistic face into the defaced data to produce nondefaced examples. The model is then trained and evaluated using a K-fold cross-validation scheme. We finally ensured the model generalizes to real nondefaced images on a public dataset that was held out. To ensure the reliability of the model against the variability introduced by the acquisition site, we included data from a diverse set of studies. A further requirement for the model was that it would be very lightweight and require minimal preprocessing, so that inference could be run on a web browser of a standard, resource-limited PC. Our contribution is three-fold: i) we propose a high-standard convolutional network model that reliably identifies nondefaced T1w MRI images of the human brain; ii) we demonstrate that very cost-effective and innovative data augmentations are possible in neuroimaging, a field characterized by data scarcity and structural biases (such as absence of nondefaced images in the case at hand); iii) the model is designed for deployment in real settings, and therefore, it is lightweight and unbiased towards a single- or a limited set of acquisition sites. Not only is the proposed tool an essential requirement to protect users and stakeholders of data-sharing initiatives from unintended release of identifiable data, but also demonstrates the potential of augmentation techniques that are increasingly gaining recognition for training machine learning models in data-limited contexts.

## 2 Materials and Methods

### 2.1 Data

All of the data are publicly available, and specific imaging parameters are found within each dataset’s distribution.

#### Training set

Our composite training dataset consisted of 980 T1w images, obtained from 36 studies publicly available at OpenNeuro.org. All of these images are distributed after the removal of facial features (i.e., defaced). All the studies in the OpenNeuro database were approved by the corresponding ethics committee before public distribution (as affirmed by the data owner upon submission).

#### Held-out evaluation dataset

To benchmark the generalizability of the model, we held-out 581 images of the publicly available IXI dataset [9, RRID:SCR 005839]. All the images in this dataset are distributed in their nondefaced form (i.e., preserving facial features).

### 2.2 Data Preprocessing

*Inpainting of synthetic faces*. The training dataset was augmented with positive-class examples (i.e., nondefaced) by inpainting a synthetic face within the defaced data (Figure 1), with AFNI’s `afni_refacer_run` [10]. After augmentation, one rater (author SB) screened every image to correct inpainting errors. Therefore, the training set’s effective sample size was duplicated to *N* =1960, with a perfect balance between defaced and nondefaced examples.

**Fig. 1:**
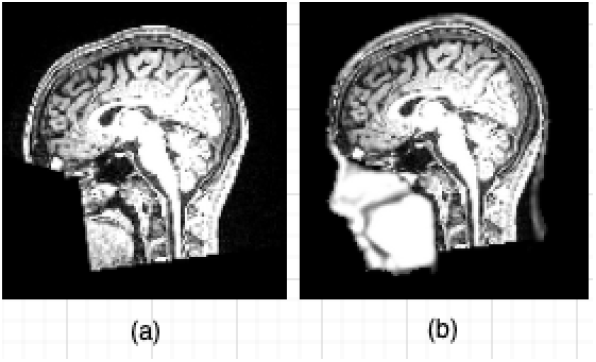
The training dataset was augmented with positive-class examples (i.e., nondefaced) by inpainting a synthetic face within the defaced data. a) A defaced image; b) corresponding nondefaced image after face in-painting.

#### Defacing of the held-out dataset

We augmented the IXI dataset with defaced examples by defacing them with *PyDeface* [6], effectively duplicating the sample size up to 1162.

#### Conformatio

All input images in NIfTI format had their data array reoriented to the RAS convention, which determines the order of the voxel axes and their direction (left-Right, posterior-Anterior, and inferior-Superior). Then, to preempt the scaling effects of the varying voxel sizes across studies, all images were resampled into a 128*×*128*×*128 grid using B-Spline cubic interpolation.

#### Intensity normalizatio

Since the data were obtained from multiple sources, the intensity range of the MRI images could also vary. To address this issue, we standardized each image’s intensity distribution to have zero mean and unit standard deviation. This was followed by a min-max normalization to ensure that voxel values were bounded between (0,1).

### 2.3 Model development and cross-validation

#### Convolutional Network Architecture

The network model (Figure 2) consists of three identical sub-models and a combined classifier at the tail. Each sub-model comprehends three “*ConvBNRelu*” layer blocks, which are a sequence of one convolutional, one batch normalization, and one non-linear activation (Rectified Linear Unit; ReLU) layers. Convolutional layers consist of *n* filters of 3×3 kernels that are used to generate feature maps. *n* starts with 8 filters in the first ConvBNRelu block and doubles every block. Batch normalization layers ensure all the features obtained as a result of convolution are normalized to have zero mean and unit variance. The ReLU layer introduces non-linearity in the function to help estimate complex decision boundaries. Each *ConvBNRelu* block is followed by a max-pooling layer. At the end of each sub-model, the features are flattened into vectors which are used by the combined classifier.

**Fig. 2:**
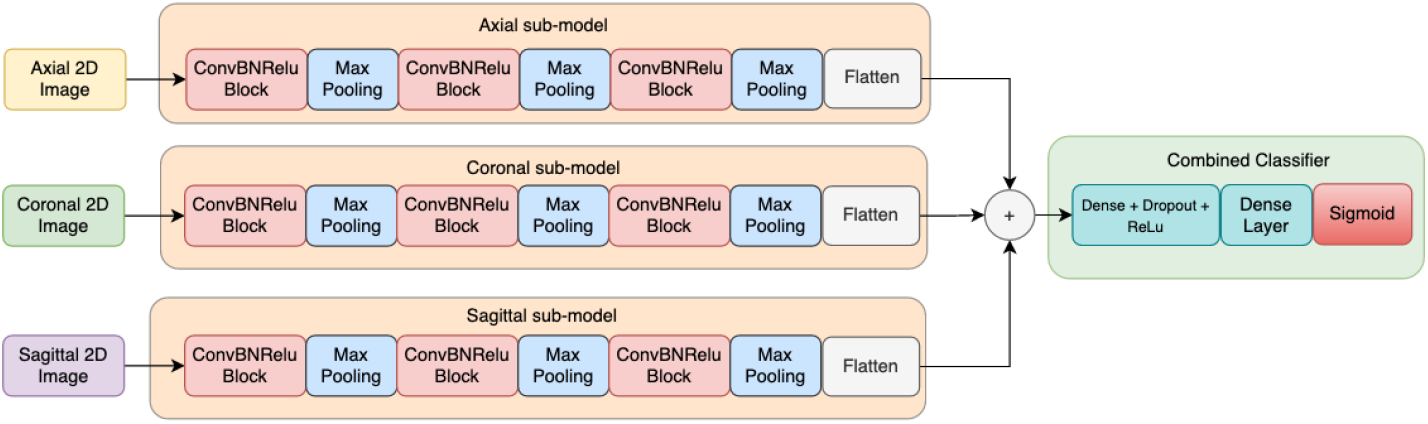
The proposed network architecture comprehends three sub-models. The network processes spatial information from all three slices (axial, coronal, sagittal) separately and then concatenates and processes the output from each sub-model to label the input image as defaced/nondefaced

The combined classifier is the final block of the network, which takes as input the flattened sum of the feature vectors from the three sub-models and outputs a real-valued probability using the sigmoid function. This probability indicates the confidence of the given input image belonging to the positive class. The proposed dense layer in the combined classifier is a combination of a fully connected and a dropout layer – fully-connected layer to map higher-dimensional features to a lower-dimensional representation while the dropouts are used to prevent over-fitting by randomly ignoring/dropping some number of layer outputs to make the training process noisier and forcing nodes within the layer to distribute the learning.

The input to each sub-model is a 2D image from the volume where each slice is acquired from the corresponding 3 planes - axial, coronal, and sagittal. For each example (i.e., 3D image), *m*=24 slices along all three axes for each submodel were randomly sampled and averaged to generate a mean slice. Averaging *m* slices is a cost-effective mechanism to ensure the input to the network sampled at locations where facial features are expected.

#### Training and optimization

Each of the sub-models is independently trained. The three sub-model blocks are trained within Step I. Upon finalization of Step I, these networks are truncated, the features are extracted and finally merged to generate the input for the combined network. In Step II, the combined classifier block is trained. After concluding the first step of training, the sub-models were frozen. Freezing the model means that the layers were excluded from any further training/weight updates. This was to ensure that only the additional layers for the combined classifier were trained in the second step of the training process. The proposed training scheme drives each sub-model to learn details of individual anatomical planes, which might not have been ensured in a single-step end-to-end training process.

We randomly sample without replacement our training set in mini-batches of 32 images (1:1 defaced:nondefaced) for each single gradient update, to ensure mini-batches contain a balanced sample of examples from sufficiently varied studies. Examples in the mini-batch are not matched, meaning that defaced examples do not correspond to their nondefaced images. As the estimated variables and labels are categorical, we used binary cross-entropy for the loss function. We used the Adam optimizer with an initial learning rate of 0.001 for training the sub-models. The Adam optimizer algorithm is regarded as being fairly robust to the choice of hyper-parameters [11] and adapts to the optimal learning rate for every variable involved after each iteration. The three sub-models were trained for 15 epochs at most, with early stopping if no improvement of the loss function is observed within a convergence window of three consecutive epochs. An epoch consists of a full cycle of *n*_train_/*n*_batch_ gradient updates. The epoch threshold value was selected to ensure that the training loss had started flattening.

In the second step of training, only the weights for the combined classifier were optimized. The initial learning rate was reduced to 5 · 10^*-*6^ to avoid the model from converging too quickly, resulting in a sub-optimal solution [12]. The classifier was trained again for up to 15 epochs with the same convergence window, using the same optimizer and the data sampling strategy.

#### Cross Validation

We overcome the potential bias that overfitting introduces when estimating performance with cross-validation (CV). We report a stratified K-fold (K=15) CV to evaluate the proposed model. The stratified sampling ensures there is no class imbalance in each fold. The average was calculated over the last trained epoch for the combined classifier model of each fold.

### Evaluation and benchmarking

#### Evaluation on a held-out dataset

Finally, we fit the model on the entire training dataset using our 2-step strategy before assessing it on the held-out set. Once the model is retrained, we run inference on the held-out dataset and report the confusion matrix (Table 1), and benchmark scores corresponding to our CV evaluation (i.e., accuracy, sensitivity, and specificity).

**Table 1:** The confusion matrix for the held-out dataset (*N* =1162) shows the very high accuracy, sensitivity, and specificity of the model on the held-out dataset (0.978/0.983/0.972, respectively).

|  |  | True class |  |  |
| --- | --- | --- | --- | --- |
|  |  | Nondefaced | Defaced |  |
| Predicted | Nondefaced | 571 | 16 | 587 |
|  | Defaced | 10 | 565 | 575 |

#### Inference time and comparison to a 3D CNN

The architecture model was trained using a GTX 1080Ti GPU, and the batch size was selected based on GPU memory capacity. We report the inference time as the average of all model predictions on the held-out dataset, without including preprocessing. Using a reference 3D-CNN, we estimate the savings in network parameters and computational requirements measured in floating-point operations per second (FLOPS). The reference 3D-CNN replaces the 2D convolutional layers within *ConVBN-Relu* blocks with corresponding 3D convolutional layers (which add an additional channel) and updating subsequent layers accordingly.

## 3 Results and Discussion

### Cross-validation estimated ∼97% sensitivity detecting inpainted datasets

Our CV evaluation of performance yielded 0.985±0.032 accuracy, 0.972±0.068 sensitivity, and 0.997±0.005 specificity averages across the K=15 folds. The narrow standard deviation around these averages add to the confidence we had on this evaluation (given the limitation imposed by not testing at all on real faces). Therefore, a training set of *N*=980 was sufficient to develop a high-sensitivity model that detects inpainted faces and confirmed the applicability of the augmentation strategy.

We interpret the results of CV as a preliminary validation that the model does not over-fit the training data. A limitation of our CV framework is that our data splitting strategy did not observe the data’s latent structure originated by the site of acquisition. In other words, our CV experiment does not confirm (or rejects) that our model is reliable on data acquired in new, unseen sites. Further analysis will be directed to evaluating our model in a leave-one-site-out folding scheme, where one entire site is held out at each CV iteration. This approach has been described in detail in our previous work [7]. The second line of improvement would be conducting a sensitivity analysis with our CV framework, modeling how the number of examples in the training set influences the final estimates of performance, determining whether the sensitivity saturates at a certain sample size.

### The model’s performance generalized well onto a held-out dataset with real (original) faces

After piloting our model and the CV evaluation, we tested that training with inpainted (synthetic) faces is a valid data augmentation to train the network on the defacing-assessment task for real T1w MRI images. Simultaneously, we give some support to the generalization of the CV results, as this second experiment is performed on the unseen IXI dataset, acquired at a new, unseen site. In simpler terms, we addressed two questions: i) whether our model would be able to identify real nondefaced images, considering that it had not been trained with even one real face; and ii) whether the model would generalize to new data (the IXI dataset). Satisfactorily, the model’s performance was high-standard and within one standard deviation of the CV figures. This result first confirmed our hypothesis that inpainted faces contain sufficient information to train a successful model for this task, which effectively overcomes the problem of nondefaced data unavailability. The second interpretation is that our stratified K-fold (K=15) CV evaluation predicted with accuracy the performance on a held-out (unseen) dataset, suggesting the model generalized to a dataset acquired at a new site. Quantitatively, the model showed very high accuracy, sensitivity, and specificity on the held-out dataset (0.978/0.983/0.972, respectively).

Although this performance is outstanding, we view the sensitivity (98.3%) as a current limitation of our work, considering the unaffordable consequences of even a single false negative (i.e., a nondefaced image that goes undetected) when openly sharing data. In addition, we cannot be certain of the features sophisticated models learn, because of their poor *interpretability*. In the case at hand, training on only synthetic faces ensures that no PII whatsoever is encoded in the model. We will continue developing this model to achieve near 100% sensitivity on the following lines: i) better understand the spatial location of the network’s attention, and the features learned by the model using Grad-CAM [13]; ii) improve the quality of the inpainted faces with more sophisticated tooling (e.g., using Abramian and Eklund’s *re-facing* network [14]); and iii) investigate potential biases analyzing patterns of false -positive and -negative inferences, in both training and held-out datasets, and construct a calibrated loss function that penalizes false negatives more than false positives.

### A lightweight model designed to perform with the highest standards with minimal resources

The average inference time on the held-out dataset was ∼12.91ms per input image. Considering that the experiments were executed on a standard GPU platform, we understand from this inference time that the model can be deployed and executed with satisfactory user-experience on a web browser of a standard PC system.

The decision to use a sub-model architecture is based on the following advantages (i) the effective augmentation of training samples as three individual sub-models are trained on slices from each plane in addition to a final combined pass instead of a single 3D image pass, (ii) less computationally intensive operations for a lightweight detector overall: 2.1M (million) total parameters and 240 MFLOPS compared to 33.6M/16,462 (parameters/MFLOPS) required to train a 3D-CNN.

### 4 Conclusion

This paper demonstrates that innovative augmentation strategies are necessary to resolve data-limited neuroimaging tasks, and key to ensure that no personally identifiable information is encoded in the model. We developed a convolutional network to detect 3D MRI images that preserve identifiable facial features. The model reached a 98.3% sensitivity on a held-out dataset, with similar specificity and accuracy scores. This model was designed to be very lightweight, with the intent of executing inference on a web browser, in resource-limited PC stations, as part of the data submission workflow of OpenNeuro.org. We will continue the improvement of the model to achieve 100% sensitivity to ensure that no identifiable information reaches unnoticed to uncertified systems and tenants.

## Data and software availability statement

All the data used in this paper are publicly available under permissive licenses. A list of the datasets retrieved from OpenNeuro.org, along with all the code this paper involves are available under an Apache 2.0 license (GitHub: poldracklab/ nondefaced-detector). We also distribute the model ready-to-use through container technology (Docker Hub: poldracklab/nondefaced-detector).

